# Molecular definition of group 1 innate lymphoid cells in the mouse uterus

**DOI:** 10.1101/330068

**Authors:** Iva Filipovic, Laura Chiossone, Paola Vacca, Russell S Hamilton, Tiziano Ingegnere, Jean-Marc Doisne, Delia A Hawkes, Maria Cristina Mingari, Andrew Sharkey, Lorenzo Moretta, Francesco Colucci

## Abstract

Determining the function of uterine lymphocytes is challenging because of the rapidly changing nature of the organ in response to sex hormones and, during pregnancy, to the invading fetal trophoblast cells. Here we provide the first genome-wide transcriptome atlas of mouse uterine group 1 innate lymphoid cells (g1 ILCs) at mid-gestation. The composition of g1 ILCs fluctuates throughout reproductive life, with Eomes^-ve^CD49a^+^ ILC1s dominating before puberty and specifically expanding in second pregnancies, when the expression of CXCR6, a marker of memory cells, is upregulated. Tissue-resident Eomes^+^CD49a^+^ NK cells (trNK), which resemble human uterine NK cells, are most abundant during early pregnancy, and showcase gene signatures of responsiveness to TGF-β, connections with trophoblast, epithelial, endothelial and smooth muscle cells, leucocytes, as well as extracellular matrix. Unexpectedly, trNK cells express genes involved in anaerobic glycolysis, lipid metabolism, iron transport, protein ubiquitination, and recognition of microbial molecular patterns. Conventional NK cells expand late in gestation and may engage in crosstalk with trNK cells involving IL-18 and IFN-γ. These results identify trNK cells as the cellular hub of uterine g1 ILCs at mid-gestation and mark CXCR6^+^ ILC1s as potential memory cells of pregnancy.

## INTRODUCTION

Most innate lymphoid cells (ILCs) reside in tissues, where they integrate the local environment and its physiology. While group 2 and 3 ILCs are well characterized across tissues in humans and mice (Nagasawa et al. 2017), the definition of group 1 (g1) ILCs is the most difficult due to their heterogeneity across tissues (Robinette et al. 2015), as illustrated by human and murine g1 ILCs in the liver (Male 2017). G1 ILCs include cytotoxic, conventional NK (cNK) cells and a host of tissue-resident ILCs in liver, uterus, spleen, gut, salivary glands and thymus, which share with cNK cells expression of surface markers, transcription factor T-bet and production of IFN-γ. Little is known, however, about the physiological role of tissue g1 ILCs throughout the body, whereas tissue ILC2s and ILC3s contribute to barrier integrity in lung and intestinal mucosa, promote tolerance of gut bacteria and regenerate lung epithelium upon viral infection (Klose & Artis 2016). G1 ILCs participate in early responses to infection through production of IFN-γ (Weizman et al. 2017; Spits et al. 2016), however conversion of cNK cells into ILC1s under the influence of TGF-β signalling undermines their anti-viral and anti-tumor responses (Gao et al. 2017; Cortez et al. 2017). Evidence also suggests g1 ILCs are involved in chronic inflammation in lung or intestine, where environmental cues drive ILC3s to convert into IFN-γ-producing ILC1s, which exacerbate pathology (Bernink et al. 2013; Bal et al. 2016). Thus, more information is available about tissue g1 ILCs in pathology than physiology (Spits et al. 2016).

Uterine ILCs contribute to optimal pregnancy outcome in mice (Boulenouar et al. 2016; Bartemes et al. 2017; Montaldo et al. 2016) and g1 ILCs are the most abundant in both human and mouse uterus (Doisne et al. 2015; Vacca et al. 2015). Among g1 ILCs, human uterine NK (uNK) cells maintain the integrity of endometrial arteries (Wilkens et al. 2013) and, during pregnancy, mediate key developmental processes and actively regulate placentation (Hanna et al. 2006) and rev. in (Moffett & Colucci 2014). For example, they modulate trophoblast invasion, reshape uterine vasculature and promote fetal growth (Hanna et al. 2006; Xiong et al. 2013; Hiby et al. 2014; Colucci & Kieckbusch 2015). Genetic epidemiology studies have shown associations of pregnancy disorders with genetic variants of Killer-cell Immunoglobulin-like Receptors (KIRs) expressed on NK and some T cells and their variable HLA-C ligands (Hiby et al. 2004; Colucci 2017). Other functions have been suggested for uterine lymphocytes, including immunological tolerance (Vacca et al. 2010), defense against pathogens (Siewiera et al. 2013; Crespo et al. 2016), and roles in pregnancy complications such as miscarriage, although the evidence for this is controversial (reviewed in (Moffett & Shreeve 2015)). Uterine ILC3s may also contribute to tissue physiology through production of IL-22, which maintains epithelial integrity (Croxatto et al. 2016). A population of immature NK cells phenotypically overlaps with ILC3s, suggesting potential plasticity between uterine g1 ILCs and ILC3s (Male et al. 2010). Mouse uNK cells are also key for uterine vascular adaptions to pregnancy (Ashkar et al. 2000). Interactions of mouse NK receptors with MHC class I molecules influence both decidual vasculature and fetal growth (Kieckbusch et al. 2014). Heterogeneity is also a feature of mouse uterine g1 ILCs, both phenotypically and functionally (Yadi et al. 2008; Zhilin Chen, Zhang, Hatta, Lima, Yadi, Colucci, Yamada & Croy 2012a; Chiossone et al. 2014), with NK cells implicated in both physiology and pathology of reproduction (Ashkar et al. 2000; Thaxton et al. 2013).

Functional heterogeneity of uterine g1 ILCs may reflect division of labour among specific subsets, and it may also result from the conversion of a subset into another under certain conditions determined by the stage of reproductive life orchestrated by sex hormones. Puberty, blastocyst implantation, placentation, parturition, and lactation are accompanied by remarkable tissue remodeling, which likely impacts on and is influenced by tissue lymphocytes. Additionally, ILC composition and function may be also marked by innate memory of pregnancy, which could contribute to the well-known better outcome of second pregnancies and their less frequent complications (Hernandez-Diaz et al. 2009).

Determining the function of uterine cell types, including ILCs, is challenging because of the changing nature of the organ and the limited access to human samples. Moreover, lack of knowledge on gene expression profiles of distinct subsets of mouse uterine g1 ILCs precludes cell type-specific gene targeting approaches in mice. Modern immunology relies on systems biology to decode cell heterogeneity and ascribe functions to discrete subsets in an attempt to define the cellular and molecular pathways of tissue physiology and pathology orchestrated by ILCs. Here we set out to begin to resolve the heterogeneity of g1 ILCs and provide the first whole-genome transcriptome atlas of mouse uterine g1 ILCs at mid-gestation.

We have previously characterized three uterine g1 ILCs (Doisne et al. 2015) and have set out here to determine their whole-genome transcriptional profile. These murine g1 ILCs are: i) Eomes^+^CD49a^+^ tissue-resident (tr)NK cells, which resemble human uNK cells; ii) Eomes^-ve^CD49a^+^ ILC1s, which may be analogous to human uterine ILC1s (Vacca et al. 2015; Montaldo et al. 2016); iii) Eomes^+^CD49a^-ve^ cNK cells, which are presumably circulating cells in both species. The results show that Eomes^+^CD49a^+^ tissue-resident uterine NK cells express genes that make them connect with most other cell types in the pregnant uterus and therefore, akin to human uNK cells, emerge as the central g1 ILC subset. Eomes^+^CD49a^-ve^ cNK cells in the uterus may support the function of tissue-resident NK cells by producing IFN-γ and responding to IL-18. Eomes^-ve^CD49a^+^ uterine ILC1s are most abundant before puberty, and CXCR6^+^ ILC1s specifically expand in the uterus, not in the liver, in second pregnancies, appearing as attractive candidates for memory cells. Determining the molecular identity and function of mouse uterine g1 ILCs may guide further work with human cells and generate opportunities for new treatments of patients with pregnancy complication and poor fecundity.

## MATERIALS AND METHODS

### Mice

C57BL/6 (B6) WT mice were purchased from Charles River UK and *Rag2*^*-/-*^*Il2rg*^*-/-*^ mice maintained in house. Eomes-GFP reporter mice were a gift of Thierry Walzer (Daussy et al. 2014), *Rorc*(*γt*)-*Cre*^TG^/R26R mice from Gé rard Eberl (Eberl & Littman 2004) and *Rosa26R-EYFP* from Ionel Sandovici. All strains were on B6 background. All animals were used at 8-12 weeks of age and were age-matched for every experiment and all time-matings. The morning of the copulation plug discovery was counted as the gestation day (gd) 0.5 for time-matings. Mice were bred, maintained and mated under pathogen-free conditions at the University of Cambridge Central Biomedical Service in accordance with the University of Cambridge Animal Welfare and Ethical Review Body and United Kingdom Home Office Regulations, as well as at the Animal Facility of the IRCCS-AOU San Martino-IST in accordance with the Italian and European Community guidelines.

### Cell isolation and functional assays

Mouse uterus, liver and spleen were processed using a protocol involving both mechanical and enzymatic processing. Finely minced tissues were digested in Accutase (Invitrogen) for 35 minutes at 37°C with gentle orbital agitation (80 rpm) (Arenas-Hernandez et al. 2015) or in Liberase DH (Roche) as described previously (Doisne et al. 2015) for functional assays. Following digestion, tissues were passed through the cell strainer (100 µm for uterus, 70 µm for liver and 40 µm for spleen) using a plunger to mechanically dissociate remaining tissues. Leucocytes were then isolated using 80%/40% Percoll gradient (GE Healthcare Life Sciences) and red blood cell lysis was performed on the resulting cell pellet using BD Pharm Lyse buffer (BD Biosciences) according to manufacturer’s instructions. Single cell suspensions obtained in this way were used for downstream analysis and assays. For *in vitro* stimulation, cells were incubated in a complete medium containing Cell Stimulation Cocktail (plus protein transport inhibitors) during a 4-hour assay, alongside Protein Transport Inhibitor Cocktail for control (both from Invitrogen).

### Flow Cytometry

Following biotin- or fluorochrome-conjugated antibodies specific for the following antigens were used: CD45 (clone 30-F11), CD3 (17A2), CD19 (1D3), CD11b (M1/70), NK1.1 (PK136), NKp46 (29A1.4), CD49a (Ha31/8), Eomes (Dan11mag), CD86 (GL1), F4/80 (BM8), B220 (RA3-6B2), MerTK (REA477), CXCR6 (SA051D1), IL-22 (1H8PWSR), NKG2D (CX5), NKG2A/C/E (20d5), CD69 (H1.2F3), CD27 (LG.3A10), CD62L (MEL-14), CD103 (2E7), Ly49H (3D10), Ly49G2 (4D11), Ly49I (YLI-90), Ly49A (YE1/48.10.6), Ly49D (4E5), Ly49E/F (CM4), MHC-II (M5/114.15.2), TruStain fcX (anti-mouse CD16/32) for Fc-receptor blocking and Fixable Viability Dye eFluor 506 for live/dead discrimination. All antibodies were purchased from Biolegend, Invitrogen (eBioscience) or BD Biosciences. For intracellular and intranuclear staining, Foxp3/Transcription Factor Staining Buffer Set (Invitrogen) was used according to manufacturer’s instructions. Uterine and liver ILC1 were sorted from Eomes-GFP reporter mice and analysed as live CD45^+^CD3^-ve^CD19^-ve^CD11b^low/-ve^NK1.1^+^NKp46^+^CD49a^+^Eomes^-ve^, uterine trNK as live CD45^+^CD3^-ve^CD19^-ve^CD11b^low/-ve^NK1.1^+^NKp46^+^CD49a^+^Eomes^+^ and uterine and liver cNK as live CD45^+^CD3^-ve^CD19^-ve^CD11b^low/-ve^NK1.1^+^NKp46^+^CD49a^-ve^Eomes^+^. Samples were acquired on a LSR Fortessa (BD Biosciences), sorted on FACSAria III instrument (BD Biosciences) and analysed using FlowJo vX software (BD Biosciences).

### RNA-sequencing

Group 1 ILCs from uterus and liver were sorted from Eomes-GFP reporter mice into the TRI Reagent (Sigma). RNA extraction was carried out by following TRI Reagent manufacturer’s instructions, followed by RNeasy Micro kit (Qiagen). cDNA was amplified using Ovation RNA-seq system V2 (NuGEN) and DNA libraries were produced using Ovation Ultralow System V2 (NuGEN) and following manufacturer’s instructions. All mRNA and cDNA quality controls and quantifications were performed using RNA 6000 Pico and High Sensitivity DNA kits (Agilent) on the 2100 Bioanalyzer instrument (Agilent). RNA sequencing was performed on Illumina Hiseq4000 at the Cancer Research UK Cambridge Institute Genomics Core. Data were aligned to GRCm38 mouse genome (Ensembl Release 84) with TopHat2 (v2.1.1, using bowtie2 v2.2.9) with a double map strategy. Alignments and QC were processed using custom ClusterFlow (v0.5dev) pipelines and assessed using MultiQC (0.9.dev0). Gene quantification was determined with HTSeq-Counts (v0.6.1p1). Additional quality control was performed with feature counts (v 1.5.0-p2), qualimap (v2.2) and preseq (v2.0.0). Differential gene expression was performed with DESeq2 package (v1.18.1, R v3.4.2) and with the same package read counts were normalised on the estimated size factors. Proportions of specific immune cell types from bulk RNA-Seq can be estimated using reference data generated from known proportions of the cell types of interest (Ziyi Chen et al. 2017). DeconRNASeq (Gong & Szustakowski 2013) was applied to take these tables of known cell proportions, defined by gene expression profiles, and used to deconvolute bulk dataset generated in this study to estimate cell proportions within each of the sequences samples.

### RT-qPCR

Whole uterus was isolated at time-points of interest, collected in RNAlater (Invitrogen) to stabilise the RNA and kept at 4°C overnight. Tissue was homogenised in the Lysing Matrix S homogenisation tubes (MP Biomedicals) on a FastPrep-24 5G Homogenizer (MP Biomedicals) using RNeasy Plus Universal Kit (Qiagen) according to manufacturer’s instruction. Isolated RNA quality control and quantification was performed using RNA 6000 Nano kit (Agilent) on the 2100 Bioanalyzer instrument (Agilent). cDNA was amplified using SuperScript VILO cDNA Synthesis kit (Invitrogen) and qPCR was performed using PowerUP SYBR Green Master Mix (Applied Biosystems), on the Quant Studio 6 Flex Real-Time PCR System (Thermo Fisher – Applied Biosystems). *Gapdh* was used as a reference gene. Data were analysed using the ^ΔΔ^Ct method. Primers were as follows: *Spp1*, 5′-TTCACTCCAATCGTCCCTAC-3′ (forward) and 5′-TTAGACTCACCGCTCTTCAT-3′ (reverse); *Ogn*, 5′-TGCTTTGTGGTCACATGGAT-3′ (forward) and 5′-GAAGCTGCACACAGCACAAT-3′ (reverse); *Ptn*, 5′-TGGAGAATGGCAGTGGAGTGT-3′ (forward) and 5′-GGCGGTATTGAGGTCACATTC-3′ (reverse) and *Gapdh*, 5′-TGCACCACCAACTGCTTAG-3′ (forward) and 5′-GGATGCAGGGATGATGTTC-3′ (reverse).

### Data and code availability

RNA-seq data are deposited in ArrayExpress (EMBL-EBI E-MTAB-6812). An open-source repository (https://github.com/CTR-BFX/2018-Filipovic-Colucci) has been created with access to the code used in this study.

### Statistical analysis

Statistical parameters and tests applied are reported in the figure legends. All statistical analyses were performed in Prism 7 (GraphPad Software) with a confidence level of 0.95. P-values above 0.05 were considered insignificant and are not indicated in the figures.

## RESULTS

### Dynamic distribution of uterine g1 ILCs during reproductive life

We have previously defined three uterine g1 ILC subsets with one closely resembling human uNK cells phenotypically (Doisne et al. 2015; Montaldo et al. 2016). These subsets are all CD45^+^CD3^-ve^ CD11b^low/-ve^ NK1.1^+^ NKp46^+^ and their surface phenotype in comparison with that of liver subsets is shown in **Figure 1A**. Here we set out to determine the distribution of g1 ILC subsets during key stages of mouse reproductive life. These were: just before puberty (3 weeks of age), during attainment of sexual maturity (5 and 8-week old), early in gestation (gd 5.5), at mid-gestation (gd 9.5), after placentation (gd13.5), late in gestation (gd17.5) and post-partum (day 1, 10 and 18). The relative abundance of different subsets at different stages was striking (**Figure 1B** and **S1A**). Before the onset of puberty and exposure to sex hormones, Eomes^-ve^ CD49^+^a ILC1s are the most abundant. During sexual maturation, ILC1s decrease, while Eomes^+^CD49a^+^ trNKs increase. Upon mating, trNK cells are the most abundant in early pregnancy, on gd 5.5. Once the placenta has been established, at gd 13.5, trNKs decrease, while Eomes^+^CD49a^-ve^ cNK cells become the most abundant. A similar landscape with cNK cells being the most abundant is observed both at day 1 and 18 post-partum, which mark respectively the beginning and the end stage of breast-feeding. Fertility is lower during lactation but not at the beginning or the end (Green, 1966). Interestingly, we observed marked differences in the distribution of g1 ILCs in the uterus of breast-feeding and non-breastfeeding females at 10 days post-partum, with breast-feeding females having increased trNK cells, similar to the distribution observed at mid-gestation (gd 9.5). These results show that the fluctuations in the three g1 ILC subsets are as dynamic as the tissue remodels. The dynamic distribution of g1 ILC subsets in the uterus at key stages of reproductive life may reflect subset-specific functions and emphasised the need for detailed characterisation of each subset.

**Figure 1.**
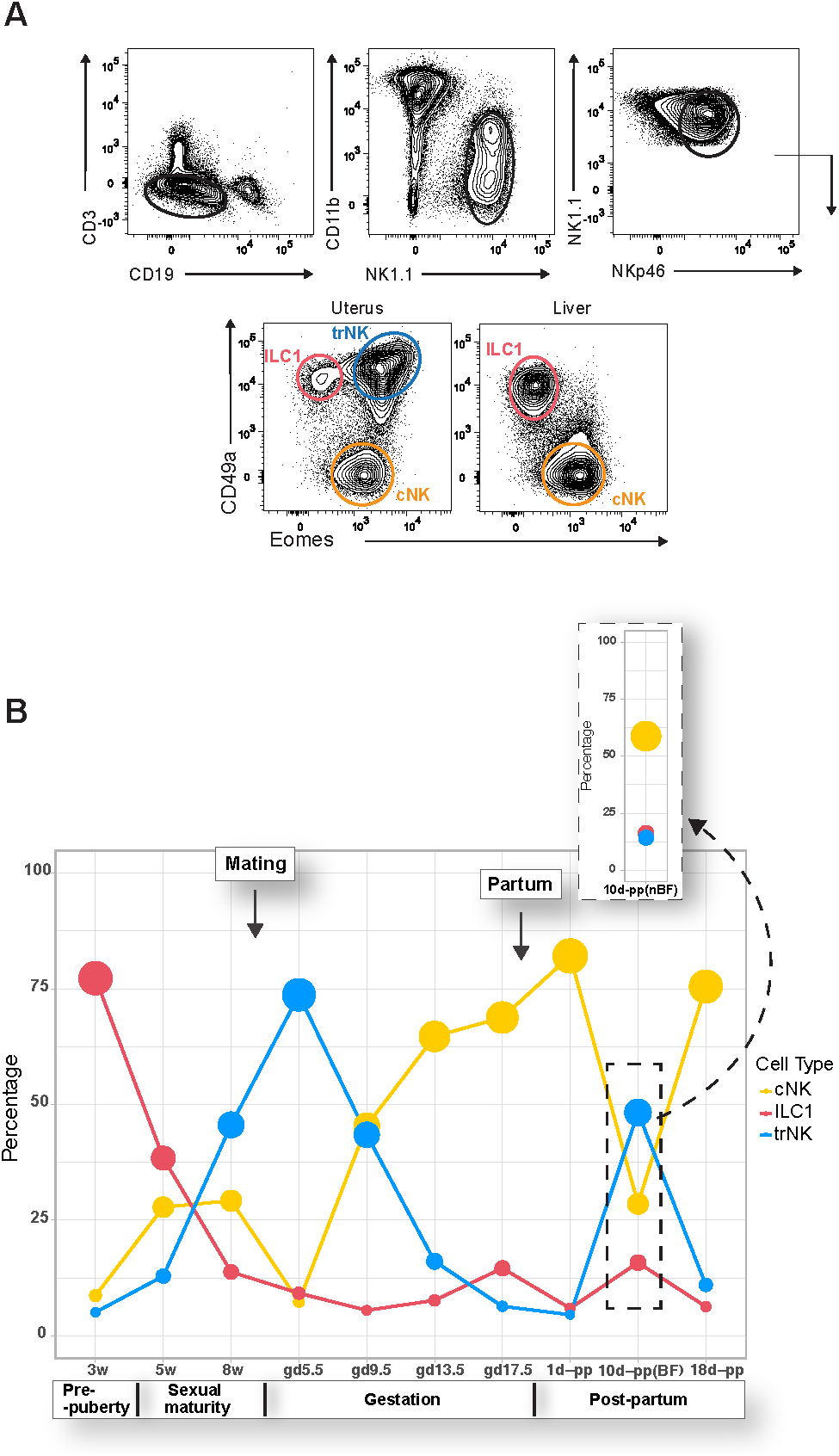
Dynamic distribution of uterine group 1 innate lymphoid cells (g1 ILCs) during reproductive life. (**A**) Gating strategy for analysis and sorting by flow cytometry, with all freshly isolated g1 ILCs at mid-gestation (gd 9.5) gated on scatter and then defined as single live CD45^+^CD3^-ve^CD19^-ve^CD11b^low/-ve^NK1.1^+^NKp46^+^; ILC1 cells defined as CD49a^+^Eomes^-ve^; tissue-resident NK (trNK) cells defined as CD49a^+^Eomes^+^ and conventional NK (cNK) cells defined as CD49a^-ve^Eomes^+^. Each panel is representative of at least a hundred independent samples. (**B**) Landscape of uterine g1 ILCs during reproductive life; numbers in plot indicate mean percentage of each individual subset of g1 ILCs as gated within live CD45^+^CD3^-ve^CD19^-ve^CD11b^low/-ve^NK1.1^+^NKp46^+^ parent population. The size of the mean data points (filled circles) is proportional to the increasing percentage of the indicated subset. Inset shows the same time-point as the dashed rectangle below but in non-breast-feeding females. Data are representative of fifteen independent experiments with three individual animals for each time-point analysed. Mating for experiments (just after animals turn 8-weeks old) and partum (gd 19.5-21.5) time-points are indicated by the arrows on the plot. w, week; d, day; gd, gestation day; pp, post-partum, BF, breast-feeding; nBF, non-breast-feeding.

### Genome-wide transcriptomes of uterine g1 ILCs at mid-gestation

Genome-wide transcriptional profiles of the three uterine g1 ILC subsets sorted from Eomes-GFP reporter mice were analysed in multiple comparisons with the two, well-established liver g1 ILCs subsets. RNA was extracted from uterine and hepatic CD45^+^ CD3^-ve^ CD11b^low/-ve^ NK1.1^+^ NKp46^+^ cells sorted based on Eomes and CD49a expression as in **Figure 1A**. This two-dimensional discrimination defines uterine and hepatic Eomes^+^CD49a^-ve^ cNK cells, uterine and hepatic Eomes^-ve^CD49a^+^ ILC1s and, unique to the uterus, Eomes^+^CD49a^+^ trNK cells. Expression of CD49a therefore marks resident cells in both tissues (Sojka et al. 2014). Hierarchical clustering of the three uterine and two hepatic subsets using the differentially expressed genes with a log2 fold change cut-off of 7.5 and P-value less than 0.05 showed that uterine and hepatic cNK cells are very similar (**Figure 2A**). Uterine trNK cells cluster close to uterine ILC1s, suggesting that the tissue micro-environment shapes the transcriptional profiles of the two uterine CD49a^+^ resident subsets. Uterine and hepatic ILC1s, however, cluster apart (**Figure 2A**), despite phenotypic similarities ((Doisne et al. 2015) and **Figure 1A**), and consistently with previously described tissue-specific gene ILC1 signatures. Principal component analysis (PCA)-based clustering confirmed the shared transcriptional profiles of uterine and hepatic cNK cells, and of uterine ILC1s and trNK cells, the PCA also highlighted differences between uterine and hepatic ILC1s (**Figure 2B**). The expression of *Tbx21* (T-bet), *Eomes, Ifng, Itga1* (CD49a), *Itga2* (CD49b), *Klrb1c* (NK1.1), and Ncr1 (NKp46) is consistent with the sorting strategy, though expression of *Tbx21, Ifng* and *Ncr1* is low in uterine ILC1s (**Figure 2C**).

**Figure 2.**
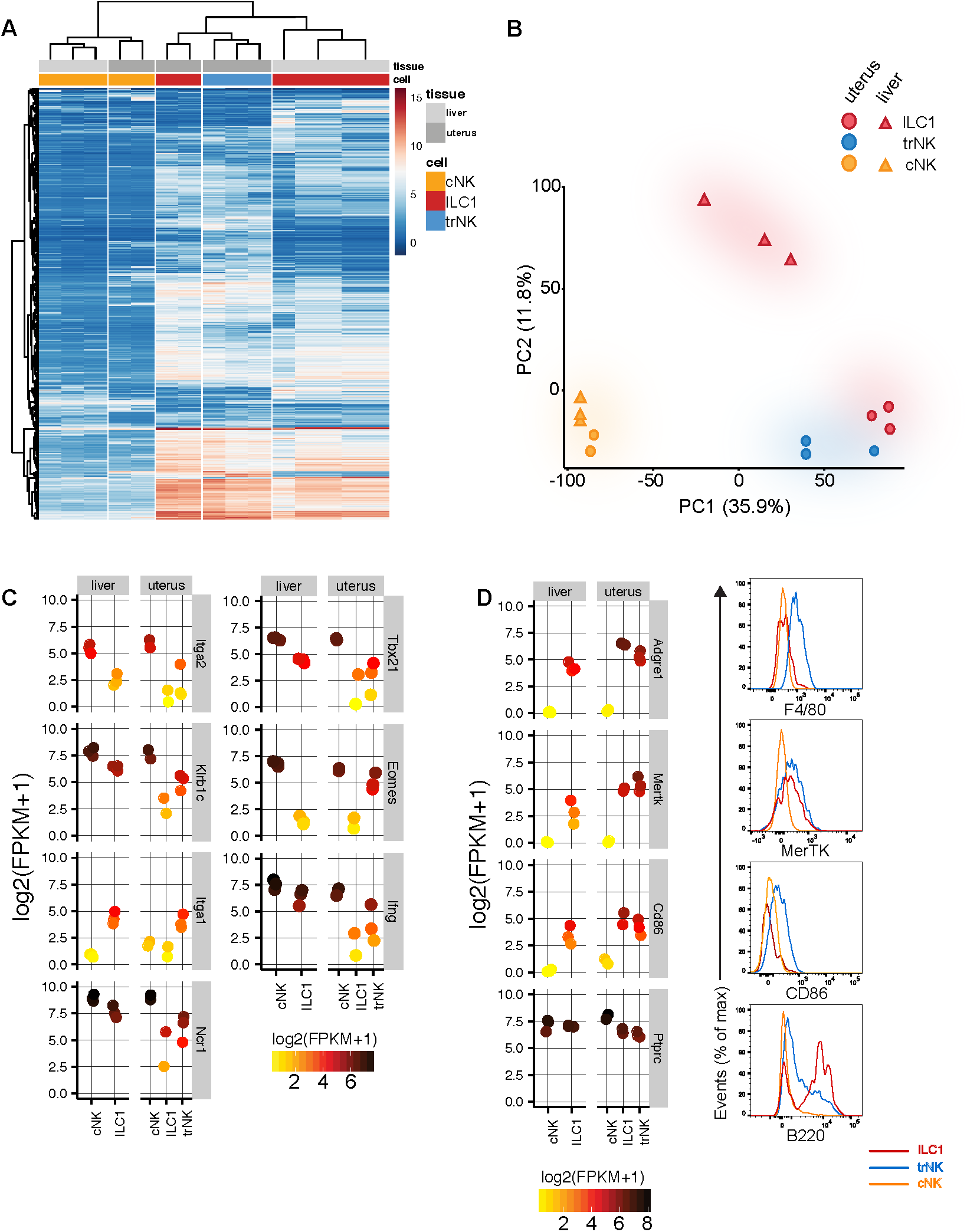
Genome-wide transcriptomes of uterine g1 ILCs at mid-gestation. (**A**) Heat map of all significant differentially expressed genes with a log2fold change >7.5 (total of 813 unique genes), selected from each of the 11 different pairwise comparisons described in the text and presented in Figure 3A. (**B**) Principal Component Analysis (PCA)-based clustering of 5 subsets of g1 ILCs: liver ILC1 and cNK and uterine ILC1, trNK and cNK cells. The first two principal components were used for clustering and to describe the variance between the subsets. (**C**) Individual gene expression plots of genes encoding proteins used for sorting g1 ILCs and additional genes used for validation of the ILC and NK-lineages. Scale represents log2(FPKM+1) transformed normalised reads. (**D**) Individual gene expression plots of genes encoding F4/80, MERTK, CD86 and B220 in uterine and liver g1 ILCs and flow cytometry validation of their protein products. Histograms show protein expression on three uterine g1 ILCs gated as gated in Figure 1A (red line-ILC1s, blue line-trNK, orange line-cNK). FPKM: fragments per kilobase of exon model per million reads mapped.

Surprisingly, genes associated with myeloid cells appeared among those defining the principal component 1 (PC1), which explained 35.9% of the variance. These included Clec7a and Clec4a3, encoding Dectin-1 and DCIR3, respectively, *Adgre1* encoding F4/80, and *Mertk* encoding MERTK, a member of the Tyro-3/Axl/Mer (TAM) family of receptor tyrosine kinases (**Figure S2A**), highly expressed in CD49a^+^ subsets compared to CD49a^-ve^ cNK cells in both uterus and liver, thus defining transcriptomic differences between resident and circulating cells in both tissues. Flow cytometry on uterine subsets showed expression of F4/80 and CD86 on trNK cells, MERTK on both trNK and ILC1s and B220 on ILC1s, with trNK expressing intermediate B220 levels (**Figure 2D**). The principal component 2 (PC2) explained 11.8% of the variance, with most genes significantly upregulated in liver ILC1s, with almost no expression in any of the other subsets, explaining in part why hepatic ILC1s do not cluster with uterine ILC1s (**Figure S2B**). These results confirm the similarities between uterine and liver cNK cells, thus aligning our results with previous work. More importantly, our results provide new data that mark the uterine resident CD49a^+^ trNK and ILC1s as unique subsets.

### Core gene signatures of uterine g1 ILCs

To compare g1 ILC transcriptomes between and among tissues, we ran 11 pairwise comparisons, the results of which are summarised in the UpSet plot (**Figure 3A**) and focused on 5 of them, as illustrated in **Figure 3B**:

I. liver (ILC1+cNK) vs uterine (cNK+trNK+ILC1);
II. liver ILC1 vs uterine (trNK+ILC1);
III. uterine (cNK vs trNK+ILC1);
IV. uterine (cNK vs trNK);
V. uterine (ILC1 vs trNK).

**Figure 3.**
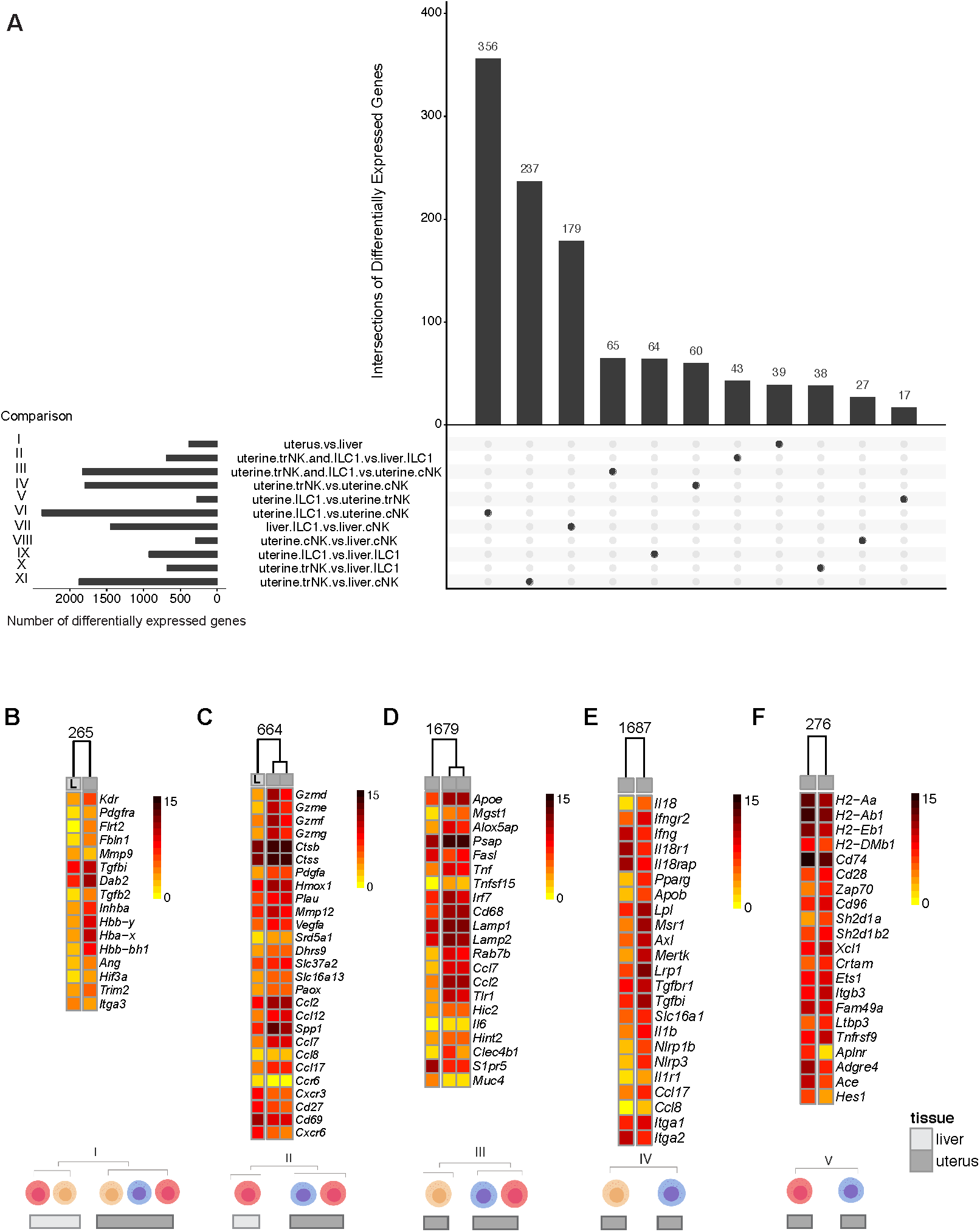
Core gene signatures of uterine g1 ILCs. **(A)** UpSetR plot displaying a summary of 11 pairwise comparisons of g1 ILCs in uterus and liver. **(B-F)** Heat maps showing selected genes for comparisons of interest (comparisons I-V in Figure 3A). Selected genes were chosen from a total list of differentially expressed genes as some representative genes from enriched pathways shown in Figure S_3_A-E. Numbers above heat maps indicate a total number of genes in a corresponding list of differentially expressed genes in that comparison: (B) comparison I: all uterine g1 ILCs vs all liver g1 ILCs. Selected genes were chosen from enriched pathways in Figure S_3_A; (C) comparison II: uterine ILC1 and trNK vs liver ILC1; Selected genes were chosen from enriched pathways shown in Figure S_3_B; (D) comparison III: uterine ILC1 and trNK vs uterine cNK cells. Selected genes were chosen from enriched pathways shown in Figure S_3_C; **(E)** comparison IV: uterine trNK vs cNK cells. Selected genes were chosen from enriched pathways shown in Figure S_3_D; **(F)** comparison V: uterine ILC1 vs trNK cells. Selected genes were chosen from enriched pathways shown in Figure S_3_E.

#### Comparison I: liver (ILC1+cNK) vs uterine (cNK+trNK+ILC1)

The comparison of transcriptomes between the three uterine and the two hepatic subsets identified 265 differentially expressed genes **(Figure 3B).** Gene Ontology (GO) analysis of this comparison for classification by biological process showed that the most highly enriched biological pathways in uterine cells relate to vascular endothelial growth factor (VEGF) signalling, including *Kdr*, encoding VEGF receptor 2 and *Pdgfra*, the alpha receptor for the platelet-derived growth factor (PDGF), but also oxygen/gas transport, including haemoglobin chains (*Hbb-Y, Hba-X, Hbb-BH1*) (**Figure S3A** and **Figure 3B**). Other significantly enriched uterine pathways included response to hypoxia and decreased oxygen levels (*Ang, Hif3α*), extracellular matrix organisation (*Flrt2, Fbln1, Mmp9, Tgfbi*), as well as several pathways relating to wound healing and the regulation of blood vessel development. TGF-β signalling genes also distinguished uterine cells, with upregulated *Dab2, Tgfb2* and *Inhba* **(Figure S3A and Figure 3B**). *Trim2* is one of the genes differentially upregulated only in the uterus and Itga3 encoding CD49c in the liver **(Figure 3B)**.

#### Comparison II: liver ILC1 vs uterine (trNK+ILC1)

The inter-tissue comparison between liver ILC1s vs uterine (trNK+ILC1), which excludes uterine and hepatic cNK cells, informs on tissue-specific profiles of CD49a^+^ resident cells **(Figure 3C).** The most highly upregulated genes in uterine CD49a^+^ cells are *Gzmd, Gzme, Gzmf, Gzmg* encoding non-cytotoxic granzymes, likely involved in tissue remodelling. Indeed, other highly expressed genes regulate extracellular matrix organisation (*Flrt2*), collagen homeostasis (*Ctsb, Ctss*), wound healing (*Hmox1, Pdgfa, Plau*), and proliferation of epithelial cells (*Mmp12, Vegfa*) (**Figures 3C** and **S3B**). Upregulated in uterine CD49a^+^ cells are also genes involved in metabolic regulation of steroids (mainly progesterone metabolism - *Dhrs9* and *Srd5a1*), ketones (*Slc37a2*) and amines (*Paox*). *Slc16a13*, which encodes an orphan monocarboxylate transporter, MCT13 (Halestrap, 2013) may be a gene specifically upregulated in uterine CD49a^+^ cells, as suggested by UpSet **(Figure 3C)**. Other upregulated genes in CD49a^+^ uterine cells are genes regulating migration of leucocytes, both myeloid cells (*Ccl2, Ccl12, Spp1*) and lymphocytes (*Ccl7, Ccl8, Ccl17*). Osteopontin-encoding *Spp1* was reported to be upregulated in uterine NK cells before (F. Wang et al. 2014). Accordingly, pregnant Rag2^-/-^Il2rg^-/-^ mice, which are devoid of all lymphocytes, and produce fetuses that are growth restricted, express less *Spp1* transcripts (**Figure S3G**). Two other growth-promoting factors, Pleiotropin-encoding *Ptn*, and Osteoglycin-encoded *Ogn* have also been implicated in NK-cell mediated fetal growth (Fu et al. 2017), however other cells produce these growth-promoting factors (Hempstock et al. 2004) and we found no significant difference in expression between wild-type and alymphoid uterus, thus excluding *Ptn and Ogn* as NK-cell and lymphocyte-specific factors driving fetal growth **(Figure S3G)**.

Expressing *Arg1, Arg2, Ecm1, and Tnfsf4* (not shown), uterine CD49a^+^ cells may regulate type-2 immunity. Unexpected was the discovery of pathways such as those regulating cellular import and phagocytosis, usually associated with antigen-presenting cells. The top enriched pathway in CD49a^+^ liver cells relative to CD49a^+^ uterine cells was negative regulation of antigen processing and presentation of peptide via MHC class II **(Figure S3B).** *H2-Oa* and *H2-Ob* (also identified by (Robinette et al. 2015) in splenic CD127^+^ ILC1s) encode H2-O, an inhibitor of MHC class II (H2-DM)-mediated antigen-loading process. Moreover, liver CD49a^+^ cells may regulate T-cell migration, with upregulated *Cxcr3* and *Ccr6.* Indeed, *Ccr6* may be exclusively expressed in liver ILC1s. Upregulated in liver cells are also *Cd27, Cd69* and *Cxcr6*, which encodes a CXCL16 receptor associated with ‘memory’ liver NK cells (Peng et al. 2013) (**Figure 3C**).

#### Comparison III: uterine (cNK vs trNK+ILC1)

We next compared the transcriptomes of the two CD49a^+^ uterine subsets with that of CD49a^-ve^ uterine cNK cells using GO-Slim for biological processes. This within-tissue comparison informs on characteristics specific to uterine resident cells. Tissue resident cells highly express genes involved in induction of cell death (*Fasl, Hvcn1, Tnf*), response to both type I interferons and IFN-γ (including chemokines *Ccl7* and very highly, *Irf7*) and proteolysis, with nearly all cathepsins significantly upregulated, alongside *C3, Plau* and *Cpq* (**Figure S3C**). This suggest that uterine resident cells may be involved in protein processing during tissue remodelling at mid-gestation. Enriched pathways are chiefly unsaturated fatty acid biosynthesis (*Mgst1*) and lipid metabolism (*Apoe, Psap*) (**Figures 3D** and **S3C**). When analysed by flow cytometry, uterine Eomes^+^ CD49a^+^ trNK cells are highly granular (Doisne et al. 2015), like their human counterpart. Consistent with this, the lysosomal transport (*CD68, Rab7b, Lamp1 and Lamp2*) is enriched in tissue resident cells. The most upregulated genes in resident CD49a^+^ cells are *Ccl7, Ccl2* and *Tlr1* (**Figure 3D**). RT-PCR analysis confirmed higher expression of *Ccl2, Ccl7*, and *Ccl12* by uterine trNK and ILC1s, compared to cNK cells (data not shown). Uterine tissue resident cells specifically upregulated the transcriptional repressor *Hic2* as well as *Il6, Hint2* and *Clec4b1*. On the other hand, highly upregulated exclusively in uterine cNK cells, as reported previously for splenic cNK cells (Robinette et al. 2015), was *S1pr5* encoding sphingosine-1-phosphate receptor 5, which regulates NK-cell egress. One of the highest fold-change upregulated genes in uterine cNK cells was *Muc4* encoding Mucin4, involved in epithelial cell integrity and homeostasis.

#### Comparison IV: uterine (cNK vs trNK)

The mutually exclusive expression of CD49a (*Itga1*) in trNK cells and CD49b (*Itga2*) in cNK cells reflected the cell-sorting strategy (**Figure 3E**). About 900 pathways were detected when we compared transcriptomes of trNK with cNK cells in the uterus (**Figure S3D**), including those involved in responses to IFN-γ and IL-18, which are differentially regulated. Upregulated in trNK cells are *Il18* and *Ifngr2*, while *Ifng, Il18r1* and *Il18rap* are upregulated in cNK cells. This suggests that trNK cells may respond to IFN-γ and produce IL-18, while cNK cells respond to IL-18 and produce IFN-γ, (**Figure 3E**). trNK cells may also respond to and produce IL-1β, through highly expressed *Il1r1, Ccl17, Ccl8* (cellular response to IL-1) and *Il1b, Nlrp1b, Nlrp3* (IL-1β production).

Among the most highly enriched pathways in trNK cells were those related to cholesterol storage and homeostasis, with genes such as *Apob, Lpl* and *Pparg* upregulated in trNK cells (**Figures 3E** and **S3D**). Interestingly, apoptotic cell clearance was among the enriched pathways, with genes like *Lrp1, Axl* and *Mertk* highly upregulated in trNK compared to cNK cells. *Lrp1* encodes CD91, a receptor of many ligands and functions. In addition to roles in apoptosis, and more widely being a haemoglobin scavenger receptor, it also binds growth factors, including TGF-β. Therefore, uterine trNK and ILC1s may be highly responsive to TGF-β, through receptors such as CD91, Tgfbr1 and Tgfbr2. In addition, *Tgfbi*, encoding transforming growth factor-beta induced protein (TGFBIp, BIGH3), was also upregulated in trNK cells. UpSet pointed to *Slc16a1*, encoding the cell exporter of lactate MCT1, to be uniquely upregulated in trNK cells, emphasizing the potential importance of anaerobic glycolysis for fueling uterine CD49a^+^ cells (Velásquez et al. 2016). Biglycan, encoded by *Bgn*, is a CD44 ligand that may be involved in the recruitment of circulating CD16^-ve^ NK cells into human endometrium (Kitaya & Yasuo 2008) and upregulated in trNK cells (**Figure S3F**). Regulators of cellular extravasation such as *Adam8, Ccl2* and *Ptafr* are upregulated in trNK, while *Itga4* and *Fam65b* are upregulated in cNK, suggesting migratory behaviour in both subsets, albeit through potentially different mechanisms **(Figure S3F)**.

*Ly86, Ly96, CD180, Sash1* are all significantly upregulated in trNK and part of highly enriched lipopolysaccharide-mediated signalling pathway. MyD88-dependent toll-like receptor (TLR) signalling was also specific to trNK cells, with most TLR-encoding genes, including *Tlr1, Tlr2, Tlr8*, and *Tlr9*, highly upregulated in trNK cells (**Figure S3F**). Lysosome organization pathway is enriched in trNK, just like complement and hemostasis regulation with genes like *C3, F7, F10* and *Ptrpj*. Also, *C3ar1* and *C5ar1*, involved in complemented receptor signaling are upregulated in trNK cells compared to cNK cells. Some of the most highly upregulated genes in trNK cells include *P2ry1* and *P2rx4*, suggesting that purinergic receptor signalling pathway is highly enriched. *Plau, Pdgfa, Sema6D, Bmpr1a, Plxna1, Itgb3* are all part of smooth muscle cell migration pathway and upregulated in trNK cells (**Figure S3F**). Other pathways enriched in trNK cells include hormone and sphingolipid catabolism, prostaglandin synthesis, detection of external biotic stimulus, regulation of nitric oxide biosynthesis, regulation of TGF-β production, and regulation of tissue remodeling. *Msr1*, encoding macrophage-scavenger receptor 1 and *Clec7a*, encoding Dectin-1 exhibited the highest fold change in trNK compared to cNK (about 300-500-fold, respectively). One of the genes upregulated almost 40-fold in trNKs is *Slc40a1*, encoding ferroportin (FPN1), which is the main exporter of iron from cells, suggesting that iron metabolism may be of importance in uterine trNK cells. Our dataset shows high expression of *Nfe2l2* (NRF2) and significantly upregulated *Nfe2l3* (NRF3) in trNK cells (Chevillard & Blank 2011). Both can bind to antioxidant stress response elements and *Nfe2l3* locus has been shown to be associated with endometriosis through genome-wide association studies (Painter et al. 2010) **(Figure S3F)**.

#### Comparison V: uterine (ILC1 vs trNK)

Comparisons I-IV highlight transcriptome profiles specific to uterine resident cells, not previously revealed for liver resident cells. Comparison V was performed to assign specific profiles to uterine ILC1s and trNK cells **(Figure 3F).** The most highly upregulated gene in uterine ILC1s compared to trNK is *Aplnr*, encoding for the adipokine apelin receptor, which is highly expressed in mouse endometrium (Pope et al. 2012). Other highly expressed genes in ILC1s include *Adgre4*, encoding the macrophage-specific F4/80 receptor, as well as *Ace*, encoding angiotensin I-converting enzyme of the renin-angiotensin system. The most highly enriched pathway in ILC1s is regulation of protein heterodimerisation activity, including *Hes1*, which interacts with the Notch pathway that is key for ILC development. Among the most highly enriched pathways detected in ILC1s is antigen processing and presentation of peptide via MHC class II, with *H2-Aa, H2-DMb1, H2-Ab1, H2-Eb1* and *CD74* highly expressed, albeit they are also expressed in trNK cells.

Enriched in trNK cells and downregulated in ILC1s are *Gzmc, Gzmd, Gzme, Gzmf and Gzmg*, assigning tissue-remodelling properties to trNK cells. Most of these granzymes were previously shown to peak in mid- to late gestation (Allen and Hamilton, 1998). Interestingly, *Xcl1* encoding lymphotactin is highly expressed by trNK cells (**Figure 3F**). XCL1 is also produced by human uNK cells and regulates trophoblast invasion (Kennedy et al. 2016a). trNK cells highly express *Cd96* encoding for the nectin-binding receptor Tactile, and *Crtam*, encoding a receptor for Nectin-like molecules (Arase et al. 2005), *Sh2d1a*, encoding SAP (Veillette et al. 2007), as well as *Sh2d1b2*, encoding EAT-2B, also a member of SAP-family of adapters (N. Wang et al. 2010). In addition, several pathways are enriched in trNK cells and related to T-cell differentiation and activation, including *CD28, Zap70*, and *Itk*. (***Figure 3F***). The trNK signature suggests also regulation of myeloid cell differentiation, through genes including *Ets1* and *Itgb3*. Other genes highly expressed in trNK cells regulate protein localization to plasma membrane, such as *Rab37, Sytl2, Ptch1, Kcnip3, Skap1, Map7* (data not shown).

### Uterine g1 ILCs express CXCR6, IL22 and some are ex-ILC3s

Although our genome-wide transcriptome analysis showed significant upregulation of *Cxcr6* in liver ILC1s (**Figure 3C**), both liver and uterine ILC1s express higher CXCR6 protein levels than trNK cells (**Figure 4A**). Because there is some overlapping between immature NK cells, ILC1s and ILC3s in other tissues (Robinette 2015), we investigated the lineage relationship among the uterine g1 ILCs using fate mapping reporter mice in which all cells that have expressed in the past, or actively express the RORγt-encoding *Rorc* gene are marked by YFP. **Figure 4B** shows presence of YFP^+^ cells among uterine g1 ILCs, which are aligned with ‘ex-ILC3s’ described in other tissues. Because the human equivalent cells reported previously as stage 3 precursor NK cells (Male et al. 2010) or NKp44^+^ ILC3s (Vacca et al. 2015) produce IL-22, we tested whether g1 ILCs can produce IL-22 and found that trNK in the mouse uterus at mid-gestation can produce IL-22, regardless of the stimulation, and to a lesser extent so can ILC1s **(Figure 4C-E).** Although Cd27 and Cd69 are both upregulated in liver CD49a^+^ cells (**Figure 3C**), CD27 and CD69 proteins are present at similar levels in both uterine and hepatic cells (**Figure 4F**). Both transcripts and Ly49 protein expression levels are slightly higher in trNK cells than in ILC1s and cNK cells within the uterus **(Figure 4F).** Despite variable levels of mRNA, all three subsets in the uterus have similar protein levels of NKG2D **(Figure 4F).** Bimodal distribution of NKG2A/C/E expression was observed for trNK cells as well. CD62L-encoding *Sell* and CD62L are both upregulated in uterine cNK, similarly to what was reported previously for splenic cNK cells (Robinette et al. 2015). CD103 is expressed by trNK cells and, to a lower level, uterine ILC1s, but not by hepatic ILC1s. However, transcripts levels of CD103-encoding *Itgae* are very low in all 5 subsets, suggesting high mRNA turnover and potential importance of CD103 in maintaining tissue-residency of g1 ILCs **(Figure 4F).** trNK cells were also more proliferative at gd 9.5, as measured by the Ki67 stain **(Figure 4F)**.

**Figure 4.**
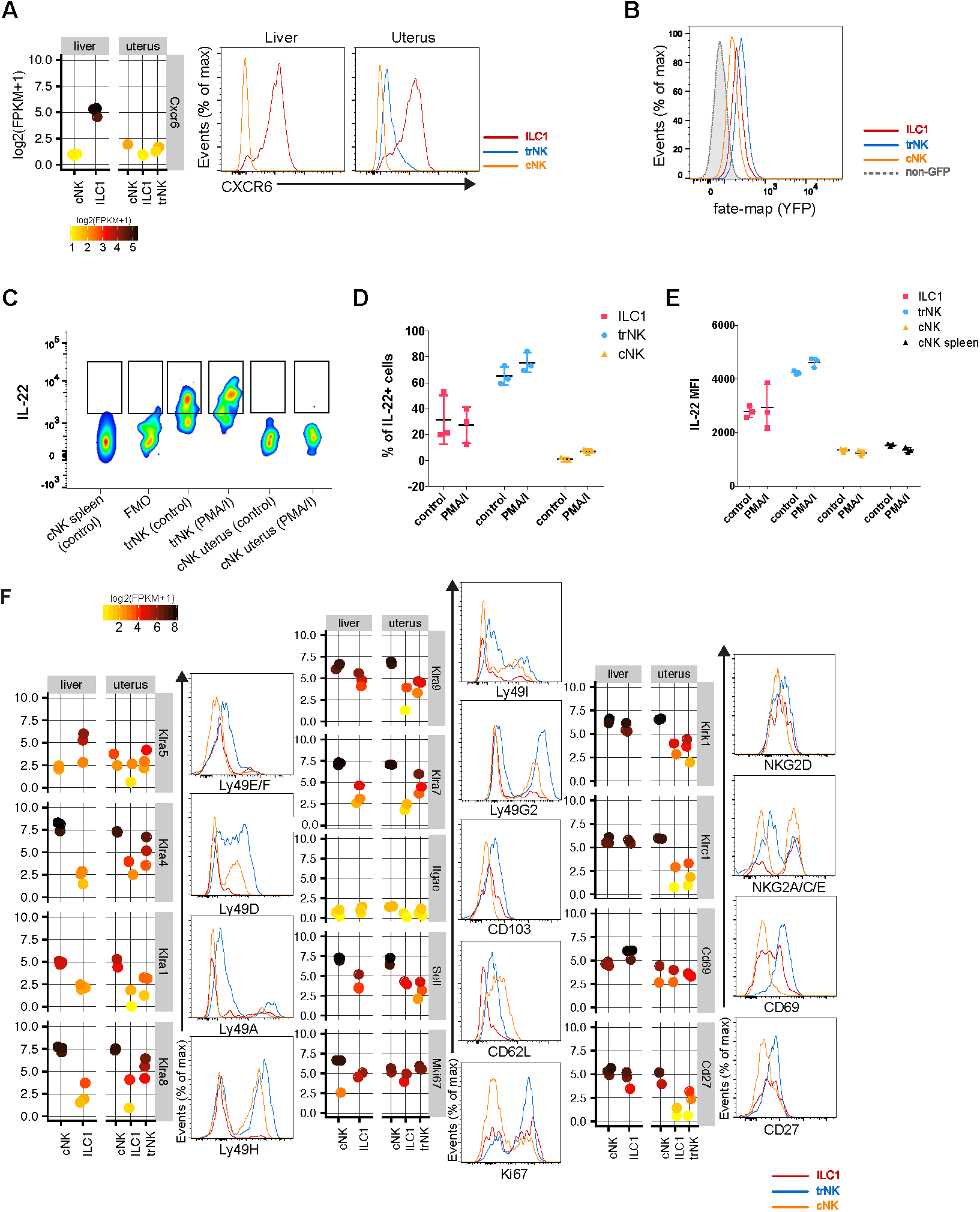
Uterine g1 ILCs express CXCR6, IL22 and some cells are ex-ILC3s. **(A)** Individual gene expression plot showing expression of *Cxcr6* in the liver and uterus. Scale represents log2(FPKM+1) transformed normalised reads. Histograms are showing CXCR6 protein expression on g1 ILCs in both organs at mid-gestation. Red line-ILC1, blue line-trNK, orange line-cNK. G1 ILCs were gated as described in Figure 1A. (**B**) Fate-mapping of g1 ILCs in the uterus of *Rorc*(*γt*)-*Cre*^TG^/R26R females. (**C**) IL-22 production by uterine trNK and cNK at gd 9.5. Control wells were incubated with medium containing protein transport inhibitors only. (**D**) Quantification of the percentage of IL-22-producing cells in the uterus at gd 9.5. Error bars represent mean±SD. (**E**) Quantification of the mean fluorescence intensity of IL-22 on g1 ILCs at gd 9.5. Error bars represent mean± SD. (**F**) Flow cytometry validation of various surface receptors on g1 ILCs in the uterus. FPKM: fragments per kilobase of exon model per million reads mapped.

### CXCR6 marks candidate memory uterine g1 ILCs of pregnancy

Adaptive, or memory NK cells have been described in humans and mice and CXCR6 is associated with memory NK and T cells in mice (Paust et al. 2010; Tse et al. 2014). It is conceivable that subsets of g1 ILCs expand in response to pregnancy-specific cues in second pregnancies and possibly contribute to the lower rate of complications in second pregnancies. We therefore compared the distribution of group 1 ILCs between first and second gestations and found no significant differences in trNK or cNK cells composition **(Figure 5A-B).** In sharp contrast, both the frequency and absolute numbers of ILC1s at mid-gestation raise about 4-5 fold in second pregnancies **(Figure 5A-B** and **Figure S5)** and ILC1s in second pregnancies upregulate CXCR6 **(Figure 5C),** suggesting these cells respond to pregnancy-specific cues and expand in second pregnancies. These results mark uterine CXCR6^+^ ILC1s as potential memory cells of pregnancy.

**Figure 5.**
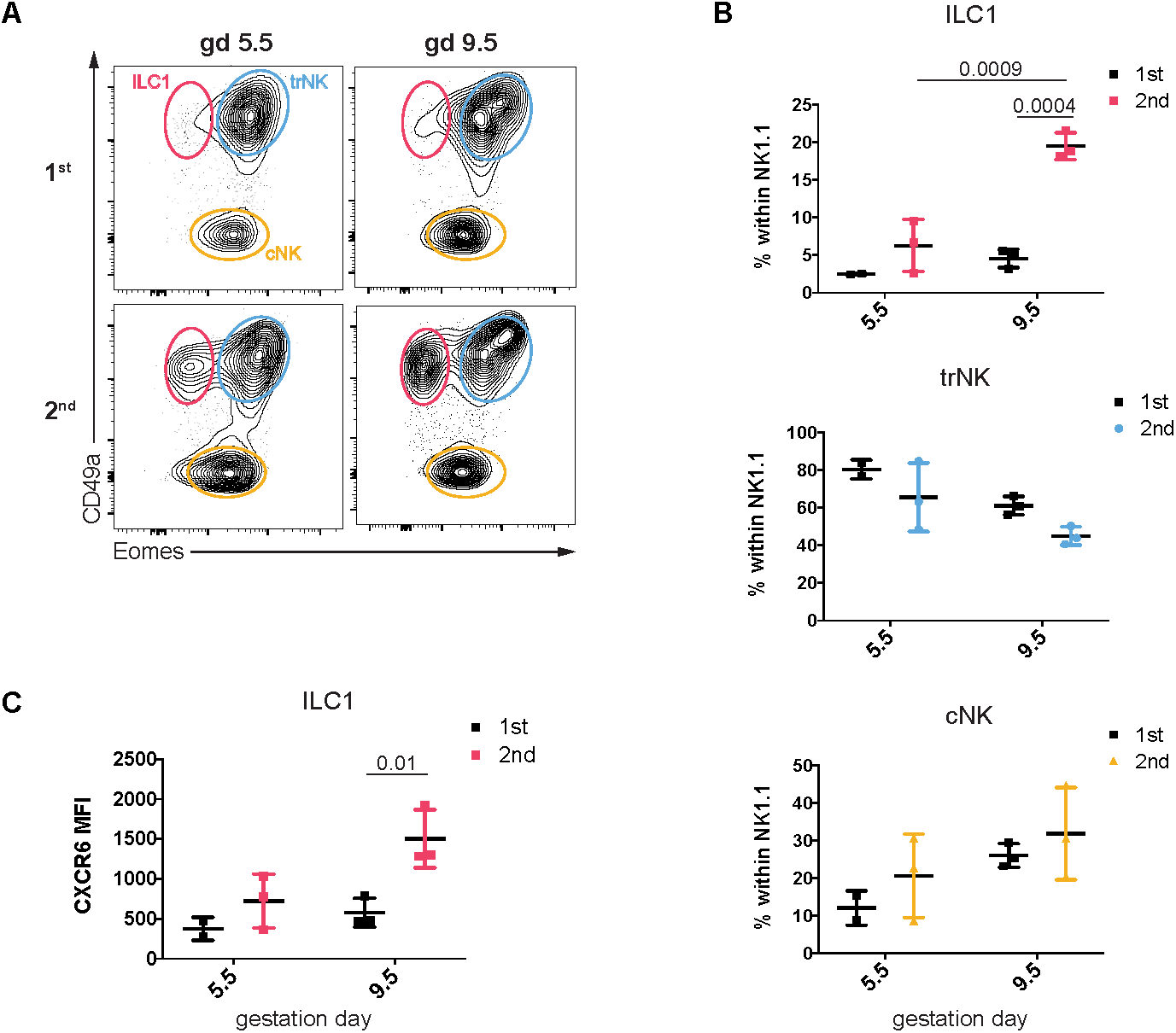
Eomes^+ve^CD49a^+^CXCR6^+^ g1 ILCs expand specifically in second gestation. (**A**) Representative plots showing distribution of g1 ILCs (gated as in Figure 1A) at gd 5.5 and gd 9.5 in first (top row) and second (bottom row) pregnancy. Data representative of 2 or 3 independent samples and one experiment. (**B**) Quantification of percentages of ILC1, trNK and cNK presented in Figure 5A in first and second pregnancy at gd 5.5 and gd 9.5. Statistical significance was evaluated by two-way ANOVA with multiple comparisons correction. Error bars represent mean±SD. (**C**) Mean fluorescence intensity of CXCR6 on ILC1s in first and second pregnancy at gd 5.5 and gd 9.5 Statistical significance was evaluated by two-way ANOVA with multiple comparisons correction. Error bars represent mean ±SD.

**Figure 6.**
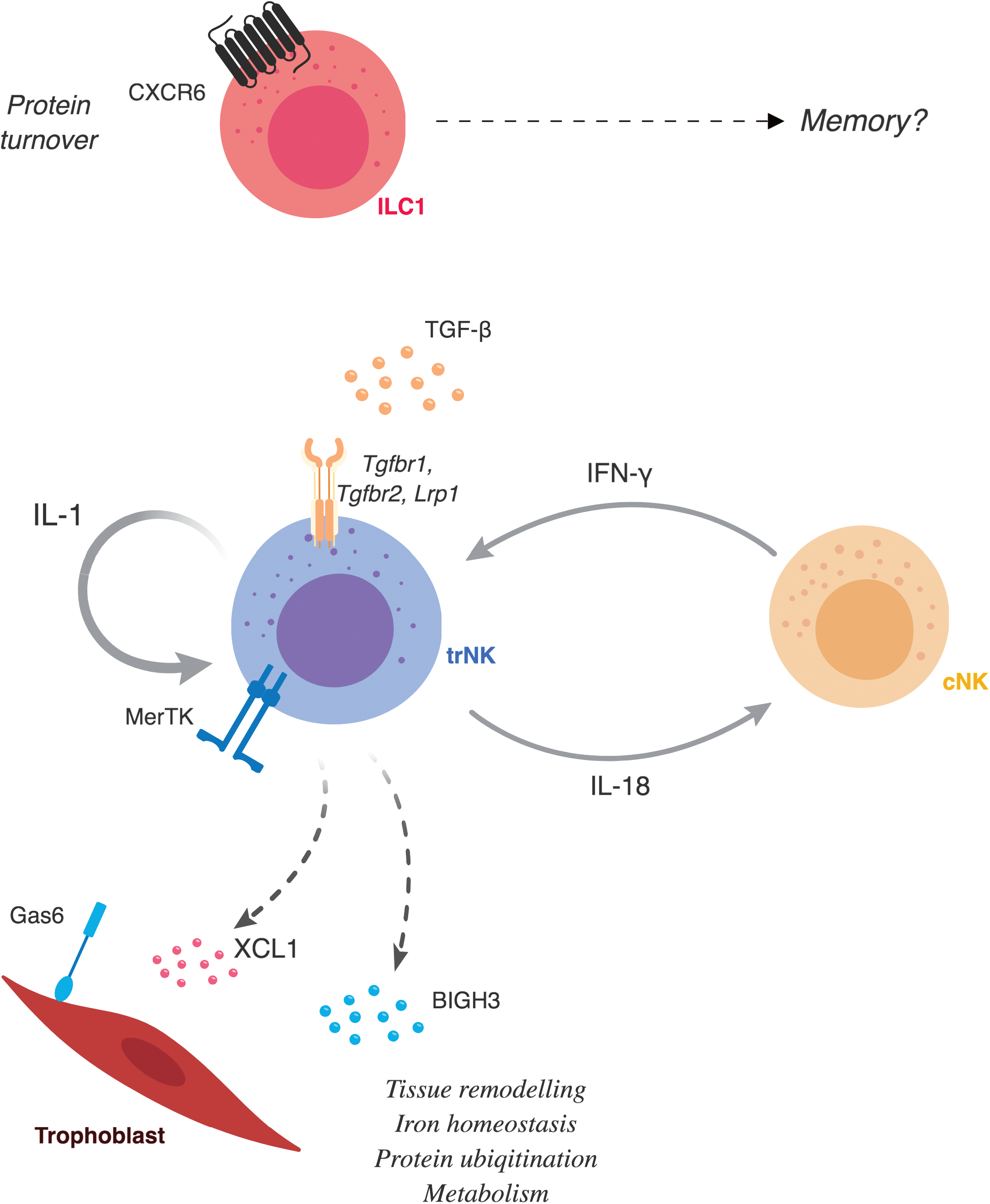
Suggested g1 ILC functions in the pregnant mouse uterus. Eomes^-ve^CD49a^+^ ILC1s are dominant before puberty and specifically expand in second pregnancies, when the expression of CXCR6 is upregulated, marking them as potential memory cells. Eomes^+^CD49a^+^ NK cells (trNK) are most abundant during early pregnancy and may well be the cellular hub of uterine g1 at mid-gestation. They indeed showcase gene signatures of responsiveness to TGF-β, connections with trophoblast, epithelial, endothelial and smooth muscle cells, leucocytes, as well as extracellular matrix. They also express genes involved in anaerobic glycolysis, lipid metabolism, iron transport and protein ubiquitination. Conventional NK cells expand late in gestation and may engage in crosstalk with trNK cells involving IL-18 and IFN-γ.

## DISCUSSION

We provide here the first whole-genome transcriptome atlas of three uterine g1 ILCs at mid-gestation in the mouse. We show clear differences in subset composition at different key stages of mouse reproductive life, demonstrating that g1 ILCs are attuned to uterine remodelling. Strikingly, we found a sharp discrimination of ILC1s and trNK cell abundance marked by puberty. ILC1s are dominant before puberty, whereas trNKs in early gestation, peaking at gd 5.5. This pattern suggests that ILC1s and trNK cells have different functions.

The picture emerging is one of trNK cells being the hub of uterine g1 ILCs, as they connect with the important cell types and factors in the pregnant uterus, including responding to TGF-β, and interacting with blood vessels, smooth muscle cells, epithelial cells of the glands, trophoblast, as well as with other leucocytes, including cNK cells that produce IFN-γ. In a cross talk similar to that of macrophages with peripheral NK cells, trNK cells may make IL-18 and stimulate cNK cells in the uterus, which in turn produce IFN-γ and stimulate trNK cells, which unexpectedly also both respond to and produce IL-1. IL-1 and IL-18 are part of the same family of cytokines, which also include the alarmin IL-33, and these are emerging as important factors in reproductive biology and pregnancy complications (Huang et al. 2016; Bartemes et al. 2017). IFN-γ is the key cytokine required for the reshaping of decidual vasculature in mice that leads to the formation of the placenta. As shown before using histological criteria for subset definitions, cNK cells are the main source of IFN-γ, while other NK-cell subsets produce angiogenic factors (Chen et al. 2012b). Our data support this division of labour among g1 ILCs, with cNK cells supporting trNK cells to engage with vasculature and other components of the pregnant uterus. Because cNK cells expand after the establishment of the placenta in the second half of gestation, it is tempting to speculate that they specialize in immune function defending the uteroplacental unit against pathogens. Future experiments will test this hypothesis.

While cNK cells in the uterus are remarkably similar to those in the liver, the core signatures of uterine resident trNK cells and ILC1s are marked by high expression of genes involved in tissue homeostasis, metabolism, genes associated with myeloid cells, cell death, interferon signalling, protein processing, and fatty acid biosynthesis, setting uterine resident trNK cells and ILC1s apart from liver resident ILC1s. The intra-tissue comparison between uterine trNK cells and cNK cells highlighted genes involved in cholesterol storage, apoptotic cell clearance, aerobic glycolysis, TLR signalling, iron transport, TGF-β signalling as well as *Axl* and *Mertk* of the Tyro-3/Axl/Mer (TAM) family of receptor tyrosine kinases. Previous work has shown that Gas6, a ligand for Mertk is expressed by trophoblast (Madeja et al. 2011). TGF-β signalling drives the transdifferentiation of cNK cells into ILC1, passing through an intermediate state that is the phenocopy of uterine trNK cells (Gao et al. 2017). Data not shown here suggests that tissue TGF-β augments in second pregnancies in the uterus and may drive a similar plasticity programme whereby cNK and trNK cells convert into ILC1s in second pregnancies, thus partly contributing to generation of candidate memory cells. In our fate mapping experiments plasticity between ex-ILC3 and g1 ILCs was evident by the reported expression of RORγt in some of the g1 ILCs. Moreover, IL-22 production by mouse trNK cells aligns them with the previously described human stage 3 uterine NK cells (Male et al. 2010), also described as NKp44^+^ ILC3s (Vacca et al. 2015).

The comparison between uterine trNK and ILC1s revealed that trNK cells contribute to tissue remodelling through non-cytotoxic granzymes, cell-cell interactions through nectin receptors CD96 and CRTAM, as well as the chemokine XCL-1 and the transforming growth factor-beta induced protein TGFBIp, also known as BIGH3. The receptor for XCL-1 is expressed in maternal myeloid cells in the human uterus and, importantly, by invading trophoblast cells too (Kennedy et al. 2016b). Therefore XCL-1, produced by both mouse and human trNK cells, emerges as a key molecule in the interactions between maternal lymphocytes and fetal cells, joining other molecular interactions, which include MHC class I receptors NKG2A, human KIR and LILRB, and murine Ly49 (reviewed in (Moffett & Colucci, 2015)). BIGH3 appears as another good candidate for the molecular interactions between resident uterine g1 ILCs as maternal endothelial, smooth muscle cells, as well as fetal trophoblast cells express BIGH3 receptors (unpublished data). With trNK and uILC1 expressing various TGF-β receptors TGF-βr1, -βr2 and CD91, it is tempting to speculate that TGF-β induces the secretion of BIGH3 by these cells, which in turn regulates downstream events essential for successful placentation, such as invasion of trophoblast, vascular remodelling, and angiogenesis. Indeed, previous studies show peak expression of uterine BIGH3 at day 4 of gestation, when trophoblast invasion starts (Uekita et al. 2003). BIGH3 has been implicated also in controlling trophoblast proliferation and invasion (Staun-Ram & Shalev 2005) as well as proliferation of trophoblast stem cells (Erlebacher et al. 2004).

The comparison between gene signatures of uterine trNK cells and ILC1s suggests that uterine ILC1s are involved in cross talk to adipocytes through the expression of the receptor for adipocytes-produced apelin. Uterine ILC1s also express the angiotensin I-converting enzyme, suggesting a potential role in regulating blood pressure, which is of paramount importance during gestation. Interestingly, trNK and ILC1s may both participate in key pathways. For example, a number of genes involved in complement, IL-1-induced plasminogen activation, as well as haemostasis appear upregulated in trNK cells, which may be involved in inflammation, its accompanying vascular response and the ensuing tissue remodelling. Genes associated with B-cells, or for protein turnover, antigen processing and presentation, as well as the macrophage-associated F4/80 marker are also upregulated in uterine ILC1s, but specific functions associated with these gene signatures remain to be determined. The role of these B-cell and myeloid markers on uterine ILC1s is unknown, and a recently described subset of NKB cells is controversial (S. Wang et al. 2017; Kerdiles et al. 2017), but might be induced in SIV and HIV infection (Manickam et al. 2018). Phenotypic and functional heterogeneity of g1 ILCs has been reported across tissues (Sojka et al. 2014; Robinette et al. 2015). The differences among g1 ILCs in different tissues may be due to different developmental pathways of separate lineages or to specific functional adaptations to different tissues. The deconvolution model we have applied here confirms that uterine g1 ILCs have expression profiles aligned with those of cell types other than NK cells and including myeloid cells. A clear limitation of the model is that there are no available ILCs defined in the dataset. Therefore, our results provide new data specific to uterine g1 ILCs at mid-gestation as a new resource for building predictive phenotype models.

Both liver and uterine ILC1s express higher levels of ‘memory’ marker CXCR6, which prompted us to look for potential changes in the subset composition in first and second pregnancies, with the idea that CXCR6^+^ ILC1s may be more abundant in second pregnancies. This was indeed the case and we suggest these cells may be associated with memory of pregnancy in the mouse and expand specifically in response to local cues. These cues remain to be determined and one possibility is that the CXCR6 ligand CXCL16, expressed also by trophoblast cells, is one driver of ILC1 expansion during pregnancy. Future work will test if and how these cells contribute to lower the rate of complications that accompany second pregnancies. For example, the hypertensive disorder of pregnancy pre-eclampsia affects 5% of primigravidae and only 1% of multigravidae (Hernandez-Diaz et al. 2009).

A number of unexpected pathways appear upregulated in tissue g1 ILC1, including those regulating iron, aerobic glycolysis, lipid storage and dectin signaling. Upregulated in trNK cells are *Fth1* and *Ftl1* encoding ferritin heavy and light chains, with *Fth1* 20-fold upregulated in trNK compared to cNK, and also one of the genes with the highest number of reads, strongly pointing to the importance of intracellular iron storage in these cells. It is therefore possible that trNKs store and export iron or, alternatively, trNK cells respond to oxidative stress induced by iron. Consistently with this possibility, *Hmox1*, which prevents inflammatory tissue injury, is upregulated in trNK cells. Along these lines, genes activated by oxidative stress and encoding enzymes that detoxify reactive intermediates through glutathione-dependent transferase and peroxidase activities, such as *Mgst1,2* are also upregulated in trNK cells. These pathways are important in the biology of DC and T cells (Rehman et al. 2013; Berod et al. 2014) and future work will establish if uterine g1 ILC require these pathways and whether their differentiation is modulated by metabolism. In a recent study, Srebp proteins and the consequent expression of genes involved in fatty-acid and cholesterol synthesis appeared essential for the metabolic reprogramming of NK cells in response to cytokine stimulation (Gardiner & Finlay 2017; Assmann et al. 2017). Interestingly, our analysis reveals that uterine, not liver resident g1 ILC upregulate *Slc16a13*, which encodes lactate and pyruvate cell exporter MCT13 (Halestrap, 2013). Warburg-like glycolysis is indeed important for decidual development (Zuo et al. 2015) and trNKs and ILC1s may provide lactate for the development of undifferentiated stromal cells in addition to other sources which function through other MCTs (MCT4). It is also tempting to hypothesise that uterine resident g1 ILCs produce lactate as a signal for arterial remodelling, in a manner analogous to how it signals for angiogenesis through acidosis in the tumour microenvironment (Romero-Garcia et al. 2016). The membrane receptor for the high-density lipoprotein cholesterol *Scarb1* is also upregulated in trNK cells, as well as pathways involved in low-density lipoprotein particle remodelling (*Abcg1, Pla2g7, Mpo*) and lipid storage (*Apoe, Soat1, Hexb, Plin2, Gm2a, Ehd1*), suggesting that lipid metabolism may be important for trNK cells. Interestingly, human SCARB1 polymorphism associates with the outcome of in vitro fertilization (Yates et al. 2011). Upregulated in g1 ILCs is also Dectin-1, an important glucan receptor for anti-fungal immunity on macrophages. Recently, peripheral NK cells were shown to recognise beta 1,3 glucan through NKp30 (Li et al. 2018), suggesting macrophage-like recognition patterns in NK cells too. Consistently with this notion, we detected a number of upregulated genes involved in pathways of microbial molecular pattern recognition. The significance of this is undetermined and future experiments of infectious challenges might shed more light on the function of these pathways in uterine g1 ILCs.

In conclusion, we provide a transcriptome atlas of uterine g1 ILCs at mid-gestation as a resource for future studies in reproductive immunology. Many genes in the UpSet lists are annotated genes that do not have a canonical name yet and include pseudogenes, long non-coding RNA and antisense transcripts. Given the general similarities of uterine trNK and ILC1s, these genes may prove to be essential in discriminating the two cell types and prove instrumental in cell-specific gene targeting. Clearly, there may be heterogeneity even within the subsets we have analysed in the uterus and it is very likely that the genes we observe in some of the subsets may be specific of smaller populations within the parent population. Future work with extensive single-cell sequencing of g1 ILCs in the uterus should capture the dynamic transcriptomic changes within cell types and stages to reveal the biology of uterine ILCs.

## ACKNOWLEDGMENTS

This work was funded by a Wellcome Trust Investigator Award 200841/Z/16/Z, the Centre for Trophoblast Research (CTR), and the Cambridge NIHR BRC Cell Phenotyping Hub to FC, the Associazione Italiana Ricerca per la Ricerca sul Cancro (AIRC)-Special Project 5×1000 no. 9962, AIRC IG 2017 Id.19920 and AIRC 2014 Id. 15283 to LM, and Ministero della Salute RF-2013, GR-2013-02356568 to PV. IF was funded by a CTR PhD fellowship. The Authors are grateful to Thierry Walzer, Gerard Eberl and Ionel Sandovici for sharing resources.

## AUTHORS CONTRIBUTION

IF designed and performed experiments, analysed data, wrote the manuscript. LC and PV designed and performed experiments, analyzed data, and revised manuscript. RSH did the computation analysis of the RNAseq data and edited manuscript. JMD designed and performed experiments, analysed data and edited manuscript. DAH and TL performed experiments. AS designed experiments, analysed data and edited manuscript. MCM and LM supervised research and revised manuscript. FC designed experiments, analysed data and wrote the manuscript.

---

**Figure S1 (Related to Figure 1). Dynamic distribution of uterine g1 ILCs during reproductive life**. (**A**) Shown are individual data points (empty rhomboids of matching colour) of the data shown in Figure 1. Means are indicated by filled circles. w, week; gd, gestation day; pp, post-partum, BF, breast-feeding; nBF, non-breast-feeding.

**Figure S2 (Related to Figure 2). First two principal components explain differences between the subsets of g1 ILCs.** (**A**) Top genes explaining variance within principal component 1 (PC1). (**B**) Top genes explaining variance within principal component 2 (PC2).

**Figure S3 (Related to Figure 3). (A-E)** Gene Ontology (GO) Analysis of selected comparisons for classifications by biological process showing selected pathways with fold enrichment: **(A)** GO for comparison I. (B) GO for comparison II. Shaded bars indicate enriched pathways in liver ILC1s. **(C)** GO-Slim for comparison III. **(D)** GO for comparison IV. **(E)** GO for comparison V. Shaded bars indicate enriched pathways in uterine ILC1s. **(F)** MA plot showing additional differentially expressed genes for comparison IV (uterine trNK vs cNK) to complement Figures 3E and S3D. **(G)** mRNA expression levels detected by RT-qPCR for *Spp1, Ptn* and *Ogn* in whole uterine tissue from either non-pregnant (virgin) B6 and *Rag2*^*-/-*^*Il2rg*^*-/-*^ mice or from B6 and *Rag2*^*-/-*^*Il2rg*^*-/-*^ dams at mid-gestation (gd 9.5). Statistical significance was evaluated by two-way ANOVA with multiple comparisons correction.

**Figure S4 (Related to Figure 4). A prediction model confirms unique features of uterine ILC1s**. (**A**) DeconRNASeq was applied to RNA-seq data generated in this study as a deconvolution method to quantitatively estimate the relative fractions of various immune cell types within gd 9.5 g1 ILCs in the liver and uterus. (**B**) Summary of the data presented in Figure S4A for each of the two subsets in liver and three subsets in the uterus.

**Figure 5S. (Related to Figure 5). Increased numbers of Eomes^-ve^CD49a^+^CXCR6^+^g1 ILCs specifically in second gestation**. The same data presented as frequency in Figure 5 are presented here as absolute cell numbers of (**A**) ILC1s, (**B**) trNK cells, (**C**) cNK cells at gd 5.5 and gd 9.5 in first (1^st^ P) and second (2^nd^ P) pregnancy. Data representative of 2 or 3 independent samples and one experiment. Statistical significance was evaluated by two-way ANOVA with multiple comparisons correction. Error bars represent mean±SD.

